# Novel inducers of the expression of multidrug efflux pumps that trigger *Pseudomonas aeruginosa* transient antibiotic resistance

**DOI:** 10.1101/655126

**Authors:** Pablo Laborda, Manuel Alcalde-Rico, Paula Blanco, José Luis Martínez, Sara Hernando-Amado

## Abstract

The study of the acquisition of antibiotic resistance (AR) has mainly focused in inherited processes, namely mutations and acquisition of AR genes. However, inducible, non-inheritable AR has received less attention and most information in this field derives from the study of antibiotics as inducers of their associated resistance mechanisms. Less is known about non-antibiotic compounds or situations that can induce AR during infection. Multidrug resistance efflux pumps are a category of AR determinants characterized by the tightly regulation of their expression. Their contribution to acquired AR relies in their overexpression. Herein we analyzed potential inducers of the expression of the chromosomally-encoded *Pseudomonas aeruginosa* clinically-relevant efflux pumps, MexCD-OprJ and MexAB-OprM. For this purpose, we developed a set of *luxCDABE*-based *P. aeruginosa* biosensor strains, which allows the high-throughput analysis of compounds able of modifying the expression of these efflux pumps. Using these strains, we analyzed a set of 240 compounds present in Biolog Phenotype Microarrays. Several inducers of the expression of the genes that encode these efflux pumps were found. The study focused in dequalinium chloride, procaine and atropine, compounds that can be found in clinical settings. Using real-time PCR, we confirmed that these compounds indeed induce the expression of *mexCD-oprJ.* In addition, *P. aeruginosa* presents lower susceptibility to ciprofloxacin (a MexCD-OprJ substrate) when dequalinium chloride, procaine or atropine are present. This work emphasizes the need of studying compounds that can trigger transient AR during antibiotic treatment, a phenotype difficult to discover using classical susceptibility tests.

## INTRODUCTION

*Pseudomonas aeruginosa* is included in the groups (ESKAPE and TOTEM) of bacteria considered to be a high risk concerning antibiotic resistance (AR) (1, 2). This nosocomial pathogen is one of the most prevalent organisms causing infections at hospitals and is the main cause of chronic lung infections in patients with cystic fibrosis and chronic obstructive pulmonary diseases (3, 4). Among the different AR mechanisms of *P. aeruginosa,* multidrug efflux pumps from the Resistance Nodulation and cell-Division (RND) family are relevant elements since they contribute to both intrinsic and acquired resistance (6–10). Among such RND efflux pumps, MexAB-OprM and MexCD-OprJ stand out as significant determinants of multidrug resistance in *P. aeruginosa* (11–13). *mexAB-oprM* is constitutively expressed under regular growing conditions, hence contributing to intrinsic resistance of *P. aeruginosa* to several antibiotics such as quinolones, macrolides, tetracycline, lincomycin, chloramphenicol, novobiocin and β-lactams (14). In addition, mexAB-oprM-overexpressing mutants have been isolated from patients (9), so that this overexpression is considered as a significant mechanism for acquiring AR under a clinic viewpoint (8). *mexCD-oprJ,* on its hand, is expressed at very low level under regular growing conditions. Thus, it does not seem to have a relevant contribution in *P. aeruginosa* intrinsic resistance (8). Nevertheless, *mexCD-oprJ* overexpression, usually achieved by loss of function mutations in the gene that encodes its local repressor, *nfxB,* confers resistance to different antibiotics such as quinolones, tetracyclines and chloramphenicol (14, 15).

It is worth mentioning that, besides being AR determinants (16, 17), efflux pumps present other physiological functions important for bacterial behavior, such as modulation of Quorum Sensing (QS) signaling (18–20), response to stress situations (21) and to host defenses (22–24), or plant/bacteria interactions (25, 26). Although the basal level of expression of each efflux pump can vary, it is well established that their expression may increase in the presence of some compounds or situations (27). In this regard, knowing which compounds are capable of triggering the expression of the genes that encode efflux pumps, and therefore to promote a transient reduction in the susceptibility to antibiotics (28, 29), a situation that is not easily detected using common susceptibility methods (8, 29), is a relevant topic.

In the current work, we addressed this issue by analyzing two relevant *P. aeruginosa* efflux pumps that present different levels of basal expression, MexAB-OprM and MexCD-OprJ. Previous studies have shown that *mexAB-oprM* expression is induced by oxidative stress (30), triclosan and pentachlorophenol (31), while envelope stress, benzalkonium chloride, chlorhexidine, tetraphenylphosphonium chloride, ethidium bromide, rhodamine 6G or antimicrobial human peptides IL-37 (32–35) are inducers of *mexCD-oprJ* expression. By the screening of a set of compounds present in Biolog Phenotype Microarrays, an approach that has been previously validated as a useful strategy to discover novel inducers of the expression of the genes that encode efflux pumps (36, 37), we have detected different molecules that induce the expression of *mexAB-oprM* and *mexCD-oprJ* in *P. aeruginosa.* For further analysis, we focused on molecules that this bacterial pathogen can potentially encounter when producing an infection, as procaine, atropine or dequalinium chloride. This work highlights the potential risk associated to the utilization of these compounds in clinical settings, as inducers of transient AR in *P. aeruginosa,* when antibiotic treatments are applied.

## MATERIALS AND METHODS

### Bacterial strains, plasmids and culture conditions

Bacterial strains and plasmids used in this work are listed in Table 1. Unless otherwise stated, all strains were grown in Lysogenic Broth, Lennox (LB) (Pronadisa) at 37°C and 250 rpm. *Escherichia coli* strains carrying the mini-CTX-*lux* (Tc^R^) or pGEM-T Easy Vector derived plasmids were grown in LB medium with 10 μg/ml of tetracycline or 100 μg/ml of ampicillin, respectively.

**Table 1.**
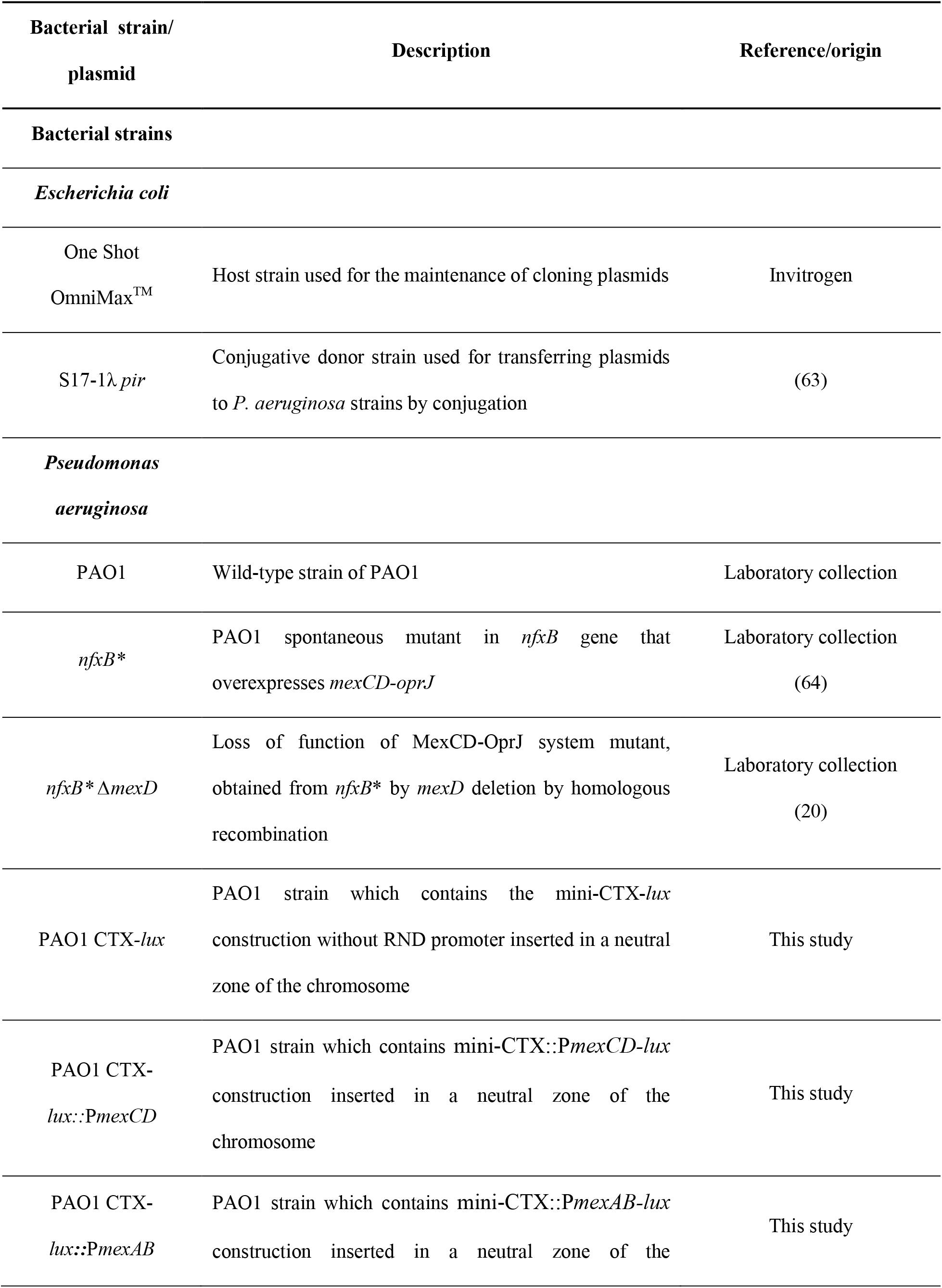
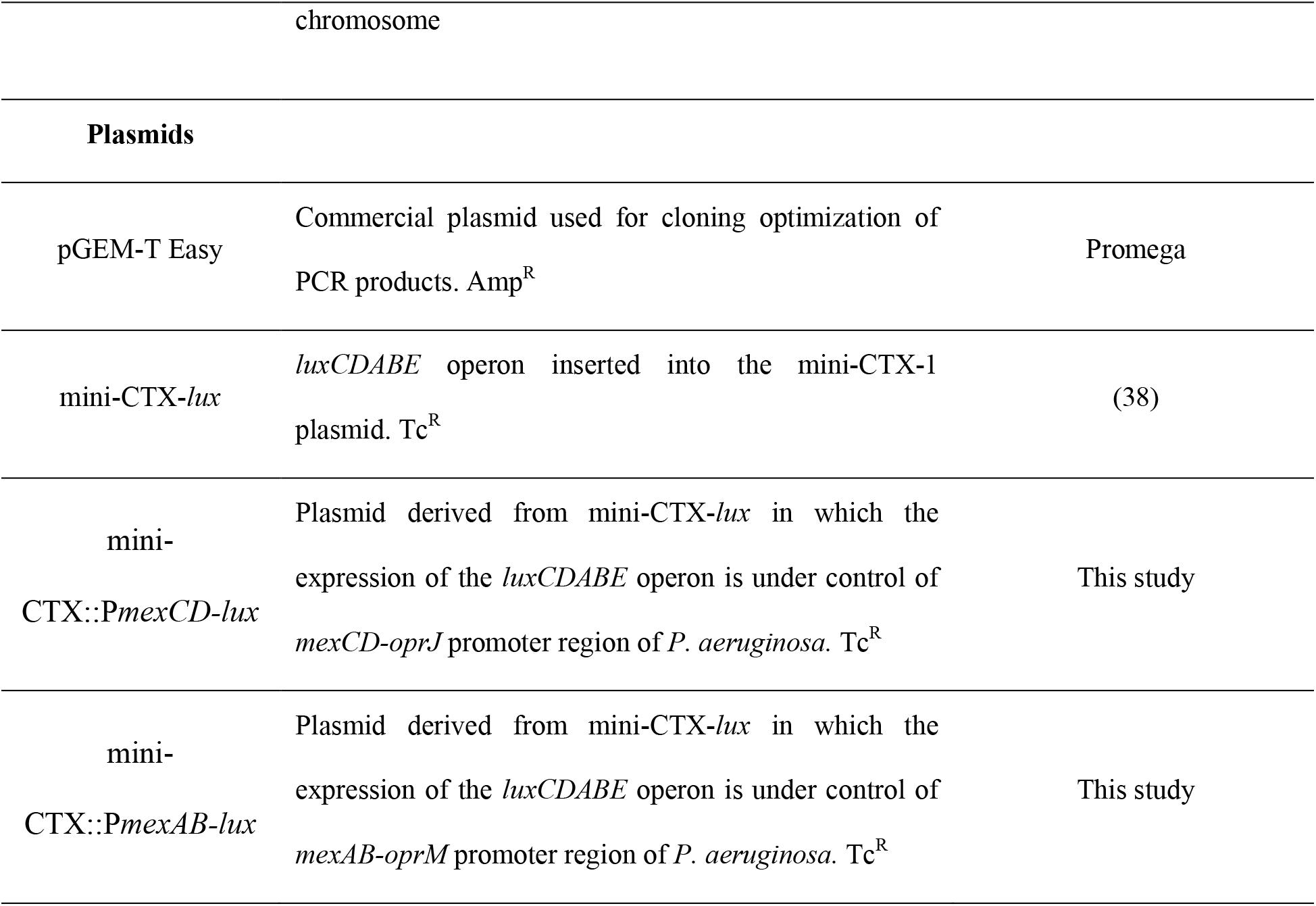
Bacterial strains and plasmids used in this work

### Construction of mini-CTX::P*mexAB-lux* and mini-CTX::P*mexCD-lux* reporter plasmids

To obtain the mini-CTX::P*mexAB-lux* and mini-CTX::P*mexCD-lux* reporter plasmids, a mini-CTX-*lux* (Tc^R^) plasmid (38) was digested with EcoRI and BamHI (New England BioLabs). The promoter region of *mexAB-oprM* operon was amplified with *EcoRI_PmexAB_Fw* and *BamHI_PmexAB_Rv* primers, while the promoter region of *mexCD-oprJ* operon was amplified with *EcoRI_PmexCD_Fw* and *BamHI_PmexCD_Rv* primers (Table 2). The products of PCR were purified from an agarose gel by using a DNA purification kit (GE Healthcare) and were cloned into the pGEM-T Easy Vector following supplier’s instructions. Afterwards, *E. coli* OmniMax^TM^ competent cells (Invitrogen) were transformed with these plasmids, which were then purified using the QIAprep Spin miniprep kit 250 (Qiagen), and digested with EcoRI and BamHI. The resulting fragments, corresponding to the efflux pump promoters, and the mini-CTX-*lux* (Tc^R^) plasmid linearized using EcoRI and BamHI, were purified from an agarose gel and used to obtain the reporter plasmids, mini-CTX::P*mexAB-lux* and mini-CTX::P*mexCD-lux*, through a ligation reaction with the T4 DNA ligase (New England BioLabs).

**Table 2.**
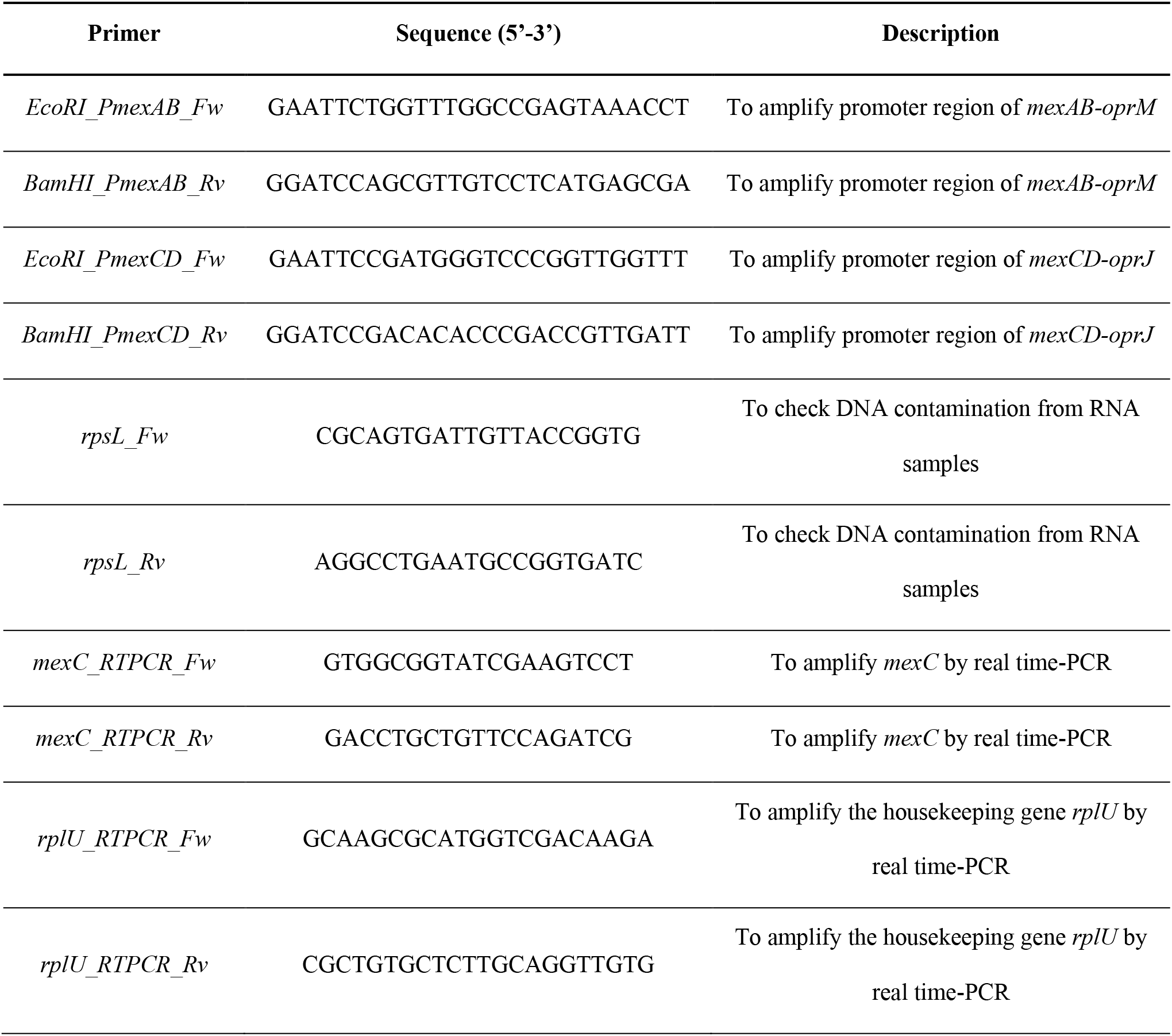
Primers used in this work

### Integration of the reporter plasmids, mini-CTX-*lux*, mini-CTX::P*mexAB-lux* and mini-CTX::P*mexCD-lux* in the chromosome of PAO1 wild type strain

The reporter plasmids, mini-CTX::P*mexAB-lux* and mini-CTX::P*mexCD-lux*, in addition to the mini-CTX-*lux*, used as control plasmid (Table 1), were introduced by transformation in *E. coli* S17-1λ *pir*. Afterwards, these constructions were independently inserted in the chromosome of *P. aeruginosa* PAO1 by conjugation, using as donor strain the *E. coli* S17-1λ*pir* harboring each of the plasmids, and following the protocol previously described (39). The *P. aeruginosa* exconjugants carrying the mini-CTX-*lux* (Tc^R^), mini-CTX::P*mexAB-lux* or mini-CTX::P*mexCD-lux* reporter constructions integrated into their chromosome were selected on petri dishes containing *Pseudomonas* Isolation Agar (PIA) (SIGMA-Aldrich) with 100 μg/ml of tetracycline. The resulting bioreporter strains are PAO1 CTX-*lux*::P*mexAB*, PAO1 CTX-*lux*::P*mexCD* and PAO1 CTX-*lux* (Table 1).

### Screening of potential inducers of *mexCD-oprJ* and *mexAB-oprM* expression

The potential ability of 240 compounds to induce *mexAB-oprM* or *mexCD-oprJ* expression was analyzed using the PAO1 CTX-*lux*::P*mexAB*, PAO1 CTX-*lux*::P*mexCD* and PAO1 CTX-*lux* strains. For that purpose, these biosensor strains were grown in from the PM11C to PM20B bacterial chemical susceptibility plates of Phenotype MicroArrays™ (BIOLOG) and both absorbance and bioluminescence emitted were monitored along time. 100 μl of LB medium were added to each well of 96-well plates, and the plates were incubated during 2 hours with agitation at room temperature to dissolve the lyophilized compounds. A volume of 10 μl of cell culture was inoculated in each well to a final OD_600nm_ of 0.01. Bacteria were grown at 37°C for 20 hours, and OD_600nm_ and luminescence was measured every 10 minutes using a Tecan Infinite 200 plate reader (Tecan).

### Normalization of the results

The ratio luminescence emitted/OD_600nm_ was calculated in those wells in which the growth rate was not impaired, and the resulting values were represented in a graphic against time. The area under the curve for each graphic was calculated using the GraphPad Prism software, giving rise to a collection of numeric values which represent the *luxCDABE* expression in PAO1 CTX-*lux*::P*mexCD*, PAO1 CTX-*lux*::P*mexAB* and PAO1 CTX-*lux* strains when growing in each specific condition. Afterwards, each numeric value obtained in each tested strain (PAO1 CTX-*lux*::P*mexAB* and PAO1 CTX-*lux*::P*mexCD*) was normalized by dividing the value corresponding to the same condition for the control strain (PAO1 CTX-*lux*).

The distribution of every normalized luminescence value obtained from each biosensor strain was represented in a box plot in order to determine the threshold from which one value will be consider as indicator of significant induction or repression. This threshold was determined as previously described (37), using the formula Q_3_ + 1.5 X IQR for the induction or Q_1_ – 1.5 X IQR for the repression, being Q_3_ the upper quartile, Q_1_ the lower quartile and IQR the interquartile range for each data set.

### Analysis of potential effectors in the expression of *mexCD-oprJ* and in the susceptibility of *P. aeruginosa* to ciprofloxacin

The growth of *P. aeruginosa* was analyzed by measuring the absorbance OD_600nm_ of bacterial cultures. 10 μl of diluted overnight bacterial cultures were added to 140 μl of LB medium with or without 0.25 μg/ml ciprofloxacin, atropine (ranging 1 to 8 mg/ml), procaine (ranging 1 to 8 mg/ml) or dequalinium chloride (ranging 1 to 500 μg/ml) in Flat white 96-well plates with optical bottom (Thermo Scientific Nunc), at a final OD_600nm_ of 0.01. To determine the effect of these potential effectors, the luminescence of either the PAO1 CTX-*lux*::P*mexCD* or the PAO1 CTX*-lux* reporter strains was measured in presence and in absence of atropine, procaine or dequalinium chloride. Measures were taken every 10 minutes during 20 or 42 hours in a plate reader (Tecan Infinite 200) at 37°C. The average of three biological replicates for each strain and condition was used to estimate the values of absorbance and luminescence.

### RNA preparation and real-time PCR

An overnight culture of *P. aeruginosa* PAO1 was used to inoculate Erlenmeyer flasks with 20 ml of LB broth to a final OD_600nm_ of 0.01. The flasks were incubated at 37°C and 250 rpm until exponential phase of growth (OD_600nm_ = 0.6) was reached. Then, the optimal concentrations of each tested compound (10 μg/ml of dequalinium chloride, 2 mg/ml of procaine and 2 mg/ml of atropine) were added to each flask and cultures were incubated for 90 minutes with shaking, as previously described in (37), to perform the induction assay; bacterial cultures without any compound or with ethanol in the case of dequalinium chloride, the solvent of this compound, were used as negative controls, and a *mexCD-oprJ* overexpressing strain (*nfxB*\*) (Table 1), grown in the absence of inducer, was used as a control of *mexCD-oprJ* overexpression. Afterwards, 10 ml of each culture were pelleted by centrifugation at 7000 rpm and 4°C for 20 minutes.

The RNA extraction from the collected cells was performed as previously described in (40). After a DNase I (Qiagen) treatment, a second treatment using DNase Turbo DNA-free (Ambion) was performed, and the presence of DNA contamination was checked by PCR using primers *rpsL_Fw* and *rpsL_Rv* (Table 2). By using the High-Capacity cDNA reverse transcription kit (Applied Biosystems), cDNA was obtained from 10 μg of RNA.

Real-time PCR was carried out with Power SYBR green PCR master mix (Applied Biosystems) in an ABI Prism 7300 real-time system (Applied Biosystems). 50 ng of cDNA were used in each reaction, except for the wells with no template that were used as negative controls. A first denaturation step at 95°C for 10 minutes was followed by 40 cycles at 95°C for 15 seconds and 60°C for 1 minute for amplification and quantification. Primers that amplify specific fragments of *mexC* were used at 400 nM (Table 2). Primers *rplU_RTPCR_Fw* and *rplU_RTPCR_Rv* were used to amplify the housekeeping gene *rplU*. All the primers used were designed with Primer3 Input software; their specificity was tested by BLAST alignment against *P. aeruginosa* genome from Pseudomonas Genome Database (http://www.pseudomonas.com/) and their efficiency was analyzed by Real-time PCR using serial dilutions of cDNA. Differences in the relative amounts of mRNA were determined according to the 2^−ΔΔCT^ method (41, 42). In all cases, the values of relative mRNA expression were determined as the average of three independent biological replicates each one containing three technical replicates.

### Determination of the susceptibility to antibiotics of *P. aeruginosa* in presence of inducers of *mexCD-oprJ* expression

Ciprofloxacin, fosfomycin, tobramycin and ceftazidime susceptibility assays were carried out using MIC-test strips (Liofilchem®) in Mueller Hinton Agar (Pronadisa) containing either 10 μg/ml of dequalinium chloride, 2 mg/ml of procaine, 2 mg/ml of atropine or without any inducer, following supplier’s instructions. Overnight bacterial cultures were normalized to an OD_600nm_ of 2.5 and a 1:1000 dilution of each culture was inoculated in the test plates and incubated at 37°C. In the case of inducers leading to small MIC changes (< 2 times), growth curves were recorded using a Tecan Infinite 200 plate reader, as previously mentioned.

## RESULTS AND DISCUSSION

### Construction and validation of reporter strains

In order to identify potential inducers of the expression of either *mexCD-oprJ* or *mexAB-oprM,* which could trigger *P. aeruginosa* non-heritable resistance to antibiotics, a set of biosensor strains of *P. aeruginosa* PAO1 containing the *mexCD-oprJ* or the *mexAB-oprM* promoter regions controlling the *luxCDABE* operon (PAO1 CTX-*lux*::P*mexCD* and PAO1 CTX-*lux*::P*mexAB* respectively) and the control strain PAO1 CTX-*lux*, were developed as described in Materials and Methods. The proper functioning of the developed reporter strains was analyzed by measuring luminescence and OD_600nm_ of each strain in the presence of known inducers, benzalkonium chloride at 10 μg/ml for *mexCD-oprJ* (34) and H_2_O_2_ at 0.136 μg/ml for *mexAB-oprM* (30). The luminescence values were normalized with respect to those obtained from the control strain PAO1 CTX-*lux*, as described in Materials and Methods. The presence of the known inducers produced an increase in luminescence emitted by the corresponding biosensor strain of 1.92-fold in the case of benzalkonium chloride, and 1.94-fold in the case of H_2_O_2_ in comparison to that produced in LB medium without inducers. These results validate the capability of these biosensor strains to detect *mexAB-oprM* and *mexCD-oprJ* inducers.

### Screening for inducers of *mexCD-oprJ* and *mexAB-oprM* expression using Biolog Phenotype Microarrays

The capability of 240 compounds, present in the PM11C to PM20B bacterial chemical susceptibility arrays of Phenotype MicroArrays™ (BIOLOG), for inducing the expression of either *mexCD-oprJ* or *mexAB-oprM* was tested. Among these compounds, there were antibiotics, heavy metals, antiseptics, fungicides, food preservatives, chelating agents, oxidative stress compounds, amino acids, synthetic organic compounds and different drugs for human and veterinary use. Four different concentrations of each compound are present in each commercial microarray plate. However, only those in which the growth of the biosensor strain was not severely impaired were considered for the analysis.

Luminescence emitted and OD_600nm_ were recorded for each biosensor strain (PAO1 CTX-*lux*::P*mexCD* and PAO1 CTX-*lux*::P*mexAB*) and these values were normalized with those obtained from the control strain (PAO1 CTX-*lux*) as described in Materials and Methods. The distribution of normalized values for each promoter was represented in a box plot (Figure 1) and an induction threshold level was calculated for each reporter strain as described in Materials and Methods. These threshold values were 1.73 and 1.53 for *mexCD-oprJ* and *mexAB-oprM* promoter, respectively. Those compounds that produced a fold change in luminescence that exceeded the calculated threshold were considered as potential inducers and are shown in Table 3. For *mexCD-oprJ,* 20 putative inducers were detected, while 10 compounds were found to increase luminescence above the threshold in the case of the *mexAB-oprM* biosensor strain.

**Figure 1.**
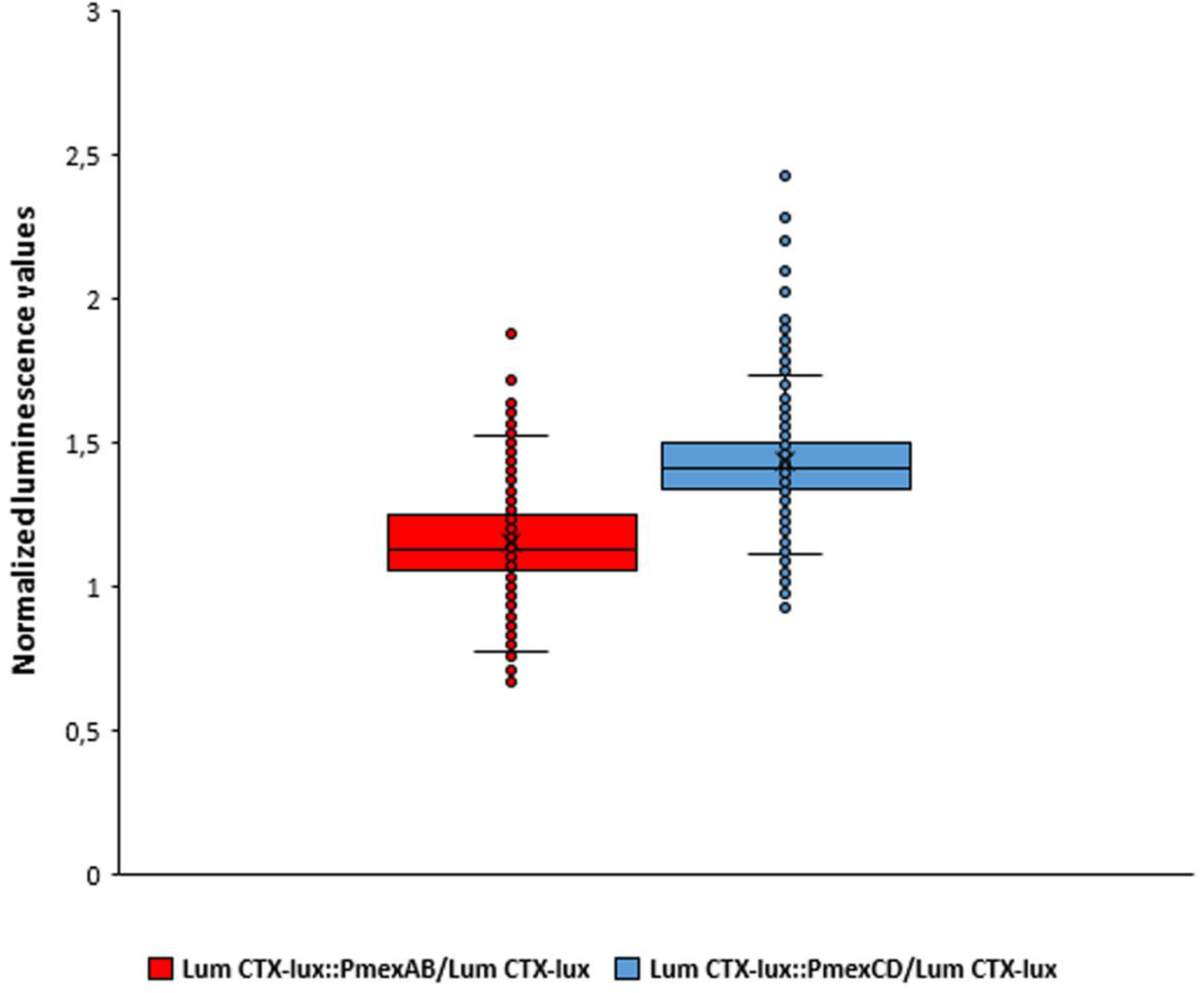
Effect of different compounds in the expression of either *mexAB-oprM* or *mexCD-oprJ*. The figure shows the normalized luminescence values produced by PAO1 CTX-*lux*::P*mexAB* and PAO1 CTX-*lux*::P*mexCD* in presence of 4 different concentrations of 240 compounds from Biolog plates. The outliers of the boxplot represent the conditions in which there was a potential overexpression or repression of the genes encoding the studied efflux pumps: those values above 1.53 for PAO1 CTX-*lux*::P*mexAB* and 1.73 for PAO1 CTX-*lux*::P*mexCD* indicate overexpression, while the values below 0.7 for PAO1 CTX-*lux*::P*mexAB* and 1.1 for PAO1 CTX-*lux*::P*mexCD* indicate repression

**Table 3.**
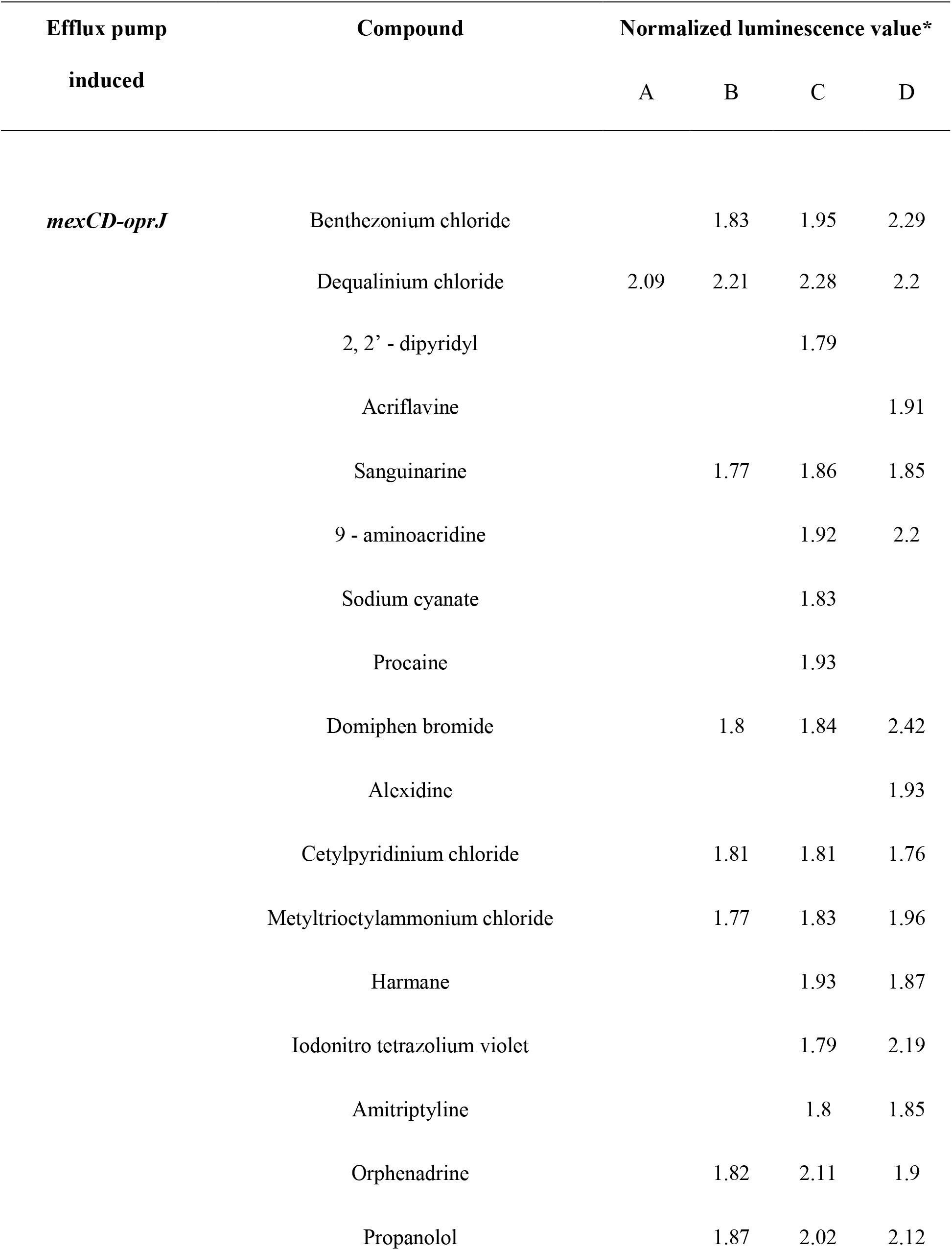
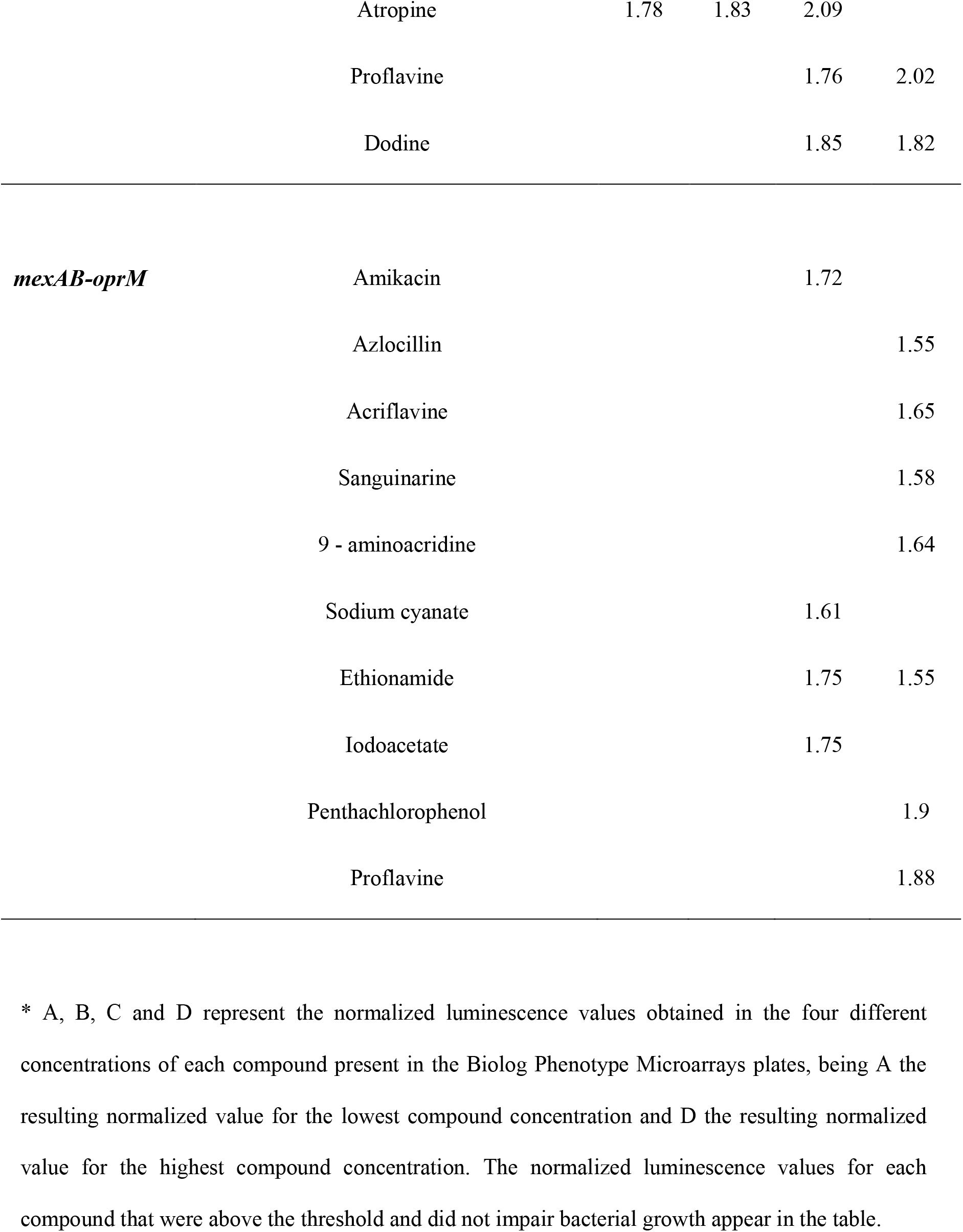
Potential inducer compounds of *mexCD-oprJ* and *mexAB-oprM* expression detected by the Biolog screening.

Among the latter, we found some compounds already known to be inducers of *mexAB-oprM* expression, which reinforce the robustness of our experimental approach. These include, pentachlorophenol, which was previously described to induce *mexAB-oprM* expression (31), or compounds that may lead to oxidative stress, which is a known inducer condition (30), such as flavins or their derivatives (acriflavine, proflavine or 9-aminoacridine) (43) or iodoacetic acid. Novel inducers of *mexAB-oprM* expression detected during the analysis include antibiotics as amikacin and azlocillin (14), sanguinarine, which is a plant-derived compound, ethionamide, which is used as an antibiotic for treating multidrug resistant *Mycobacterium tuberculosis* (44) and sodium cyanate, a neurotoxic compound implicated in neurodegenerative disorders in populations subsisting on the cyanogenic plant cassava (45) (see Table 3).

Concerning MexCD-OprJ, an efflux pump known to be induced by membrane-damaging agents (32, 33), novel putative inducers of its expression were found. Among them, disinfectants (benthezonium chloride, dequalinium chloride, domiphen bromide, alexidine and cetylpyridinium chloride), the chelating agent 2, 2’-dipyridyl, sodium cyanate, methyltrioctylammonium chloride, iodonitro tetrazolium violet, flavin derivatives (acriflavine, proflavine and 9-aminoacridine), the anesthetic agent procaine, the antidepressant drug amitriptyline, the antihistaminic agent orphenadrine, the β-blocker propranolol, the fungicide dodine and some plant-derived compounds (atropine, harmane and sanguinarine), were identified (see Table 3).

Regarding plant-derived compounds, it is worth mentioning that some of them, as the flavonoids, play important physiological functions in plants as well as in plant-bacteria interactions (46), and it is known that the flavonoid-responsive efflux pump MexAB-OprM of *Pseudomonas syringae* is required for the efficient bacterial colonization of tomato plants (47). Since *P. aeruginosa* is widely distributed in different natural habitats, including plants (48), it seems possible that these efflux pumps could have a role in colonizing such habitats (25, 26).

In addition to the detected inducers of *mexCD-oprJ* and *mexAB-oprM* expression, some compounds (Table 4) were found to reduce the expression of these efflux pumps below the lower threshold (1.1 for *mexCD-oprJ* and 0.7 for *mexAB-oprM)* for each of both promoters calculated as described in Materials and Methods. For *mexCD-oprJ,* those compounds were the antibiotics chloramphenicol, spectinomycin and spiramycin, and the plant derived compounds nordihydroguaiaretic acid and gallic acid (see Table 4). For *mexAB-oprM,* there were antibiotics such as hygromycin and josamycin, chromium chloride, glycine hydroxamate and protamine sulfate (see Table 4). However, these compounds did not seem to be strong inhibitors, since the normalized luminescence values measured in presence of them were close to those of the threshold values and were not further studied.

**Table 4.**
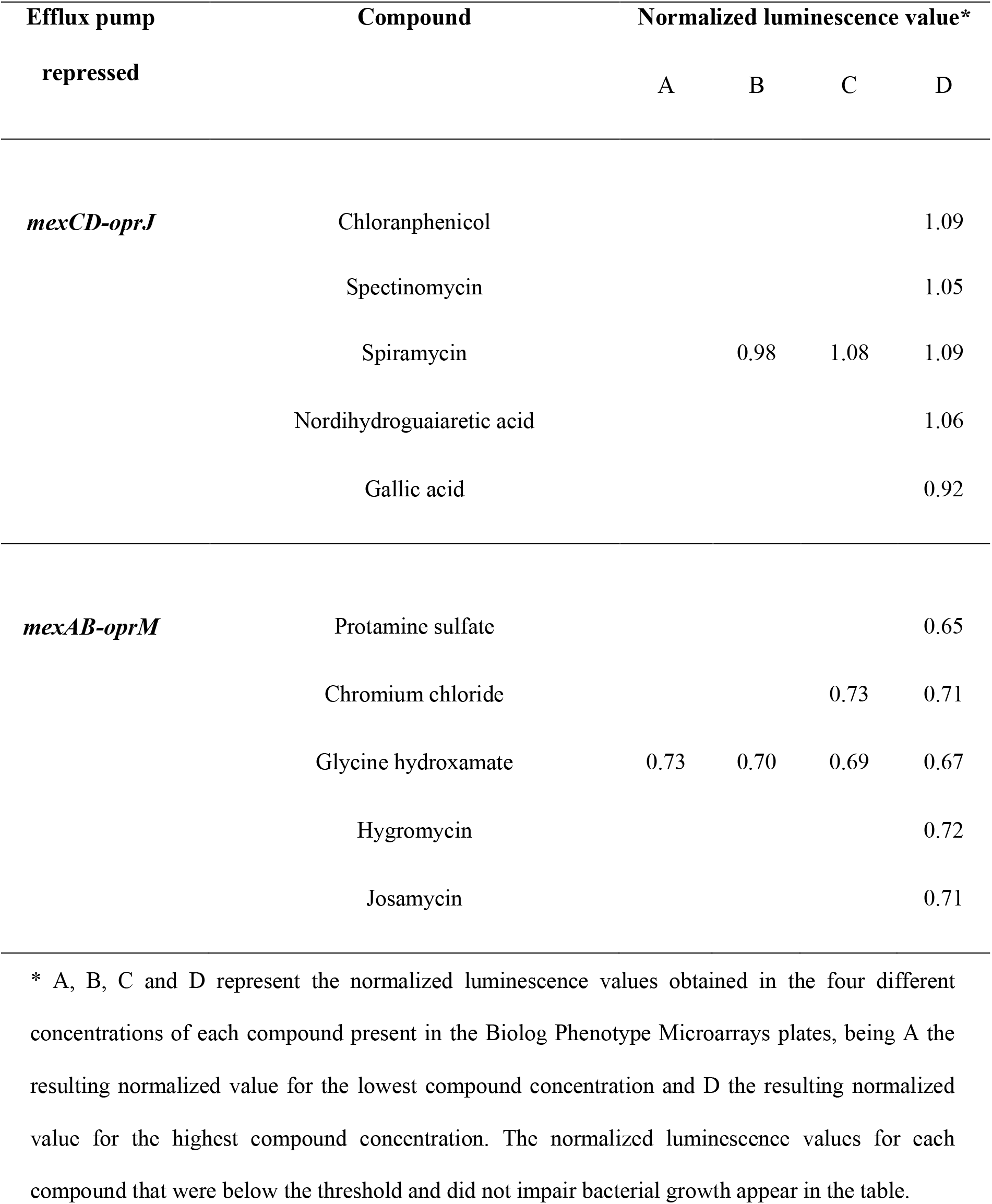
Potential inhibitors of *mexAB-oprM* and *mexCD-oprJ* expression detected with the Biolog screening.

Among the potential inducers, dequalinium chloride, procaine and atropine, which seemed to induce the expression of *mexCD-oprJ*, were chosen for a deeper analysis because they are used in human therapy and hence *P. aeruginosa* can grow in their presence when causing infections.

### Inducers of expression of *mexCD-oprJ* with relevance in clinical settings

Dequalinium chloride is used as a disinfectant (49) and presents structural similarities to the known inducers benzalkonium chloride and chlorhexidine (34), which promote the *mexCD-oprJ* expression through the induction of an envelope-stress condition (32, 33). Further, several compounds found as inducers in the analysis are similar to them in structure and/or biological activity (benthezonium chloride, cetylpyridinium chloride, domiphen bromide or alexidine) and are also used in clinical practice. This finding supports the potential relationship between antibiotics and disinfectants resistance, as well as the possible induction of AR by disinfectants of common use. In particular, the induction of *mexCD-oprJ* by dequalinium chloride could have clinical relevance, as this compound forms part of several antiseptic and disinfection procedures (50) and its use has been considered for the treatment for promyelocytic leukemia (51–54).

Procaine is used as a local anesthetic agent (55) in some minor surgeries or in burn injuries, tissues that *P. aeruginosa* frequently colonize (56). Moreover, it has been studied for the treatment of HIV patients (phase 2 clinical trial) (57). Therefore, the induction of *mexCD-oprJ* by procaine should be taken into consideration in these patients since they are very susceptible to *P. aeruginosa* infections (58). Finally, atropine is considered as an essential drug for preoperative medication and sedation by the World Health Organization (44), so it may have an effect on *P. aeruginosa* susceptibility to antibiotics when this microorganism infects surgical patients.

Since concentrations of each compound in the Biolog plates are unknown, the *mexCD-oprJ* reporter strain was grown during 20 hours in a range of concentrations in order to select one at which an increase in luminescence is observed but bacterial growth is not compromised. These concentrations are 10 μg/ml for dequalinium chloride, 2 mg/ml for procaine and 2 mg/ml for atropine, and were the concentrations used for further studies. Luminescence measurements in the presence of inducers at these optimal concentrations were recorded and normalized as previously described. In all cases, the presence of the tested compounds increased the luminescence produced by the biosensor strain (Figure 2), further supporting they induce the expression of *mexCD-oprJ.*

**Figure 2.**
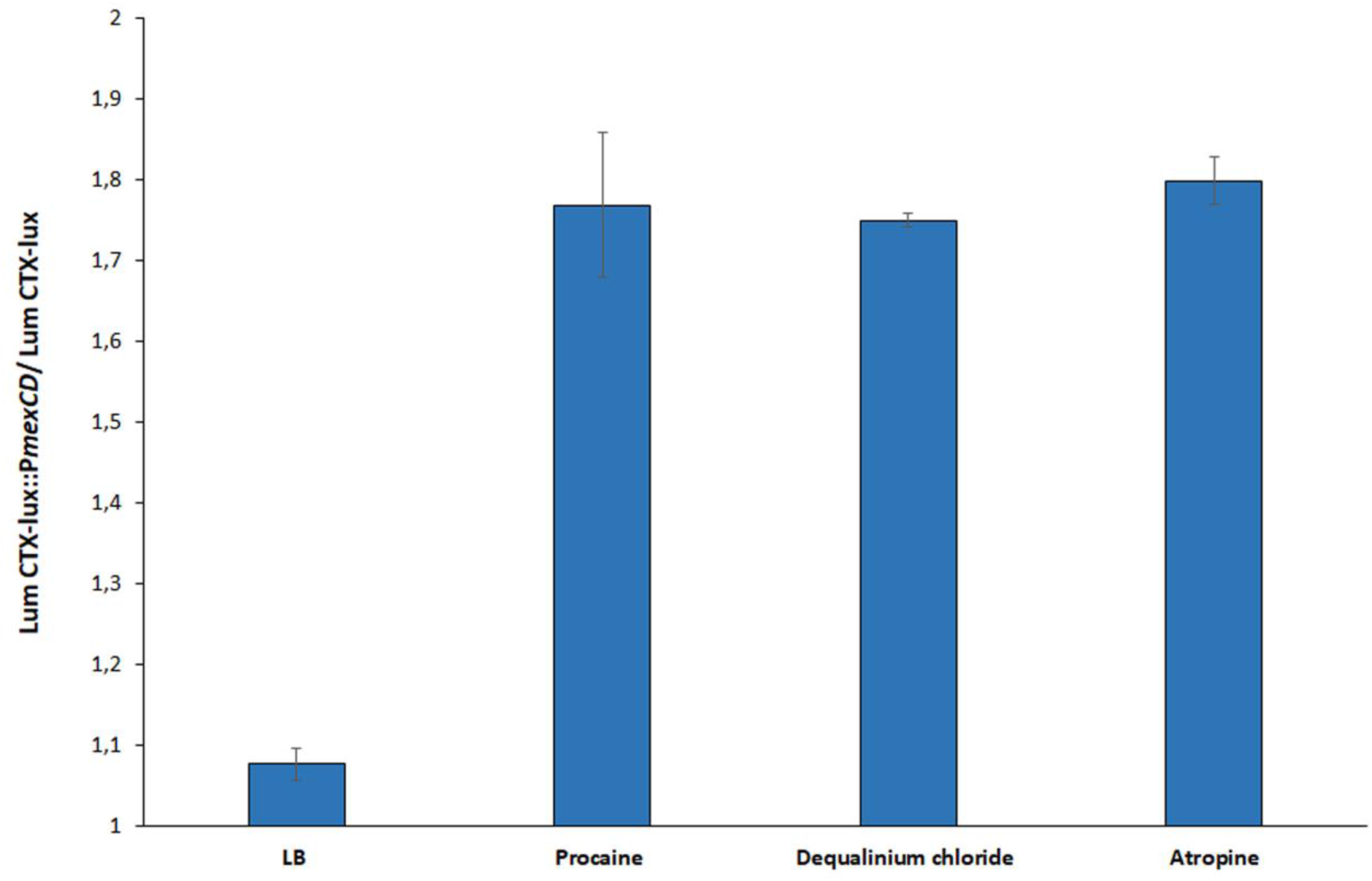
Effect of dequalinium chloride, procaine and atropine in the expression of *mexCD-oprJ*. The Figure shows the normalized luminescence values produced by the reporter strain PAO1 CTX-*lux*::P*mexCD* in presence of 10 μg/ml of dequalinium chloride, 2 mg/ml of procaine and 2 mg/ml of atropine. Luminescence was normalized to that produced by PAO1 CTX-*lux* in presence of inducer. As shown, expression of *mexCD-oprJ* is induced by the three tested compounds. Error bars represent standard deviation of three independent replicates.

### Dequalinium chloride, procaine and atropine induce *mexCD-oprJ* expression and transient antibiotic resistance

In order to further confirm the inducing capacity of the compounds found in the screening, the expression level of *mexCD-oprJ* was quantified by real-time PCR in the wild type *P. aeruginosa* PAO1, grown in the presence or in the absence of the cognate inducer compounds. The strain overexpressing *mexCD-oprJ (nfxB*)* (Table 1) was used as efflux pump-overexpressing control strain. The expression of *mexCD-oprJ* increased by 54-fold in the presence of dequalinium chloride, 25-fold in the presence of procaine and 16-fold in the presence of atropine (Figure 3). These results confirm that the expression of *mexCD-oprJ* is induced by the compounds selected from the Biolog screening.

**Figure 3.**
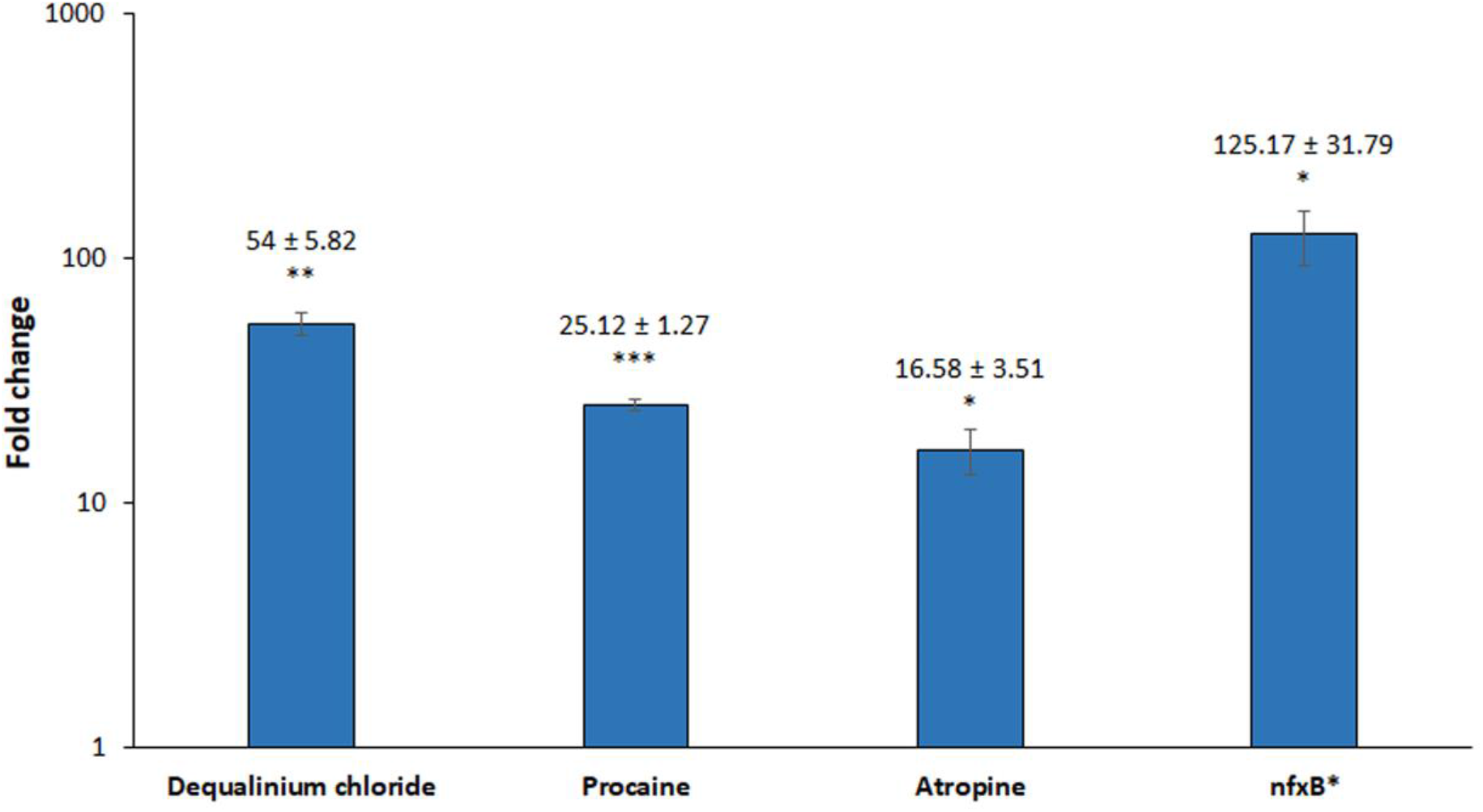
Analysis of *mexCD-oprJ* expression by real time-PCR in the presence of inducers. *mexC* expression was measured by real-time PCR after 90 minutes of incubation with 10 μg/ml of dequalinium chloride, 2 mg/ml of procaine, 2 mg/ml of atropine or without inducer. Fold changes were calculated regarding the expression in *P. aeruginosa* PAO1 untreated. The *nfxB\** strain grown in the absence of any inducer was used as a control of overexpression. Each represented value is the average of three biological replicates. As shown, expression of *mexCD-oprJ* is induced by the three tested compounds. Statistically significant differences regarding PAO1 untreated were calculated with *t*-test for paired samples assuming equal variances: *p < 0.05; **p < 0.005; ***p < 0.0005.

The *P. aeruginosa* PAO1 MIC of ciprofloxacin, one of the MexCD-OprJ substrates, was analyzed in the presence and in the absence of the aforementioned inducers in order to determine the effect of such induction on the susceptibility of *P. aeruginosa* to this antibiotic. A strain that overexpresses *mexCD-oprJ (nfxB*)* and a mexD-defective mutant (*nfB*\*Δ*mexD*) (Table 1) were used as controls. MIC of ciprofloxacin was higher in the presence of the inducers (Table 5). Moreover, the presence of each inducer did not affect the MIC of ciprofloxacin for *nfxB\** and *nfxB*ΔmexD,* confirming that the observed phenotype was caused specifically by *mexCD-oprJ* induction. In the case of atropine, which leads to lower increase of ciprofloxacin MIC (2-fold), growth curves of PAO1 in presence of ciprofloxacin, with or without this inducer, were also analyzed (Figure 4). The results confirm that atropine leads to a transient reduction of *P. aeruginosa* ciprofloxacin susceptibility.

**Figure 4.**
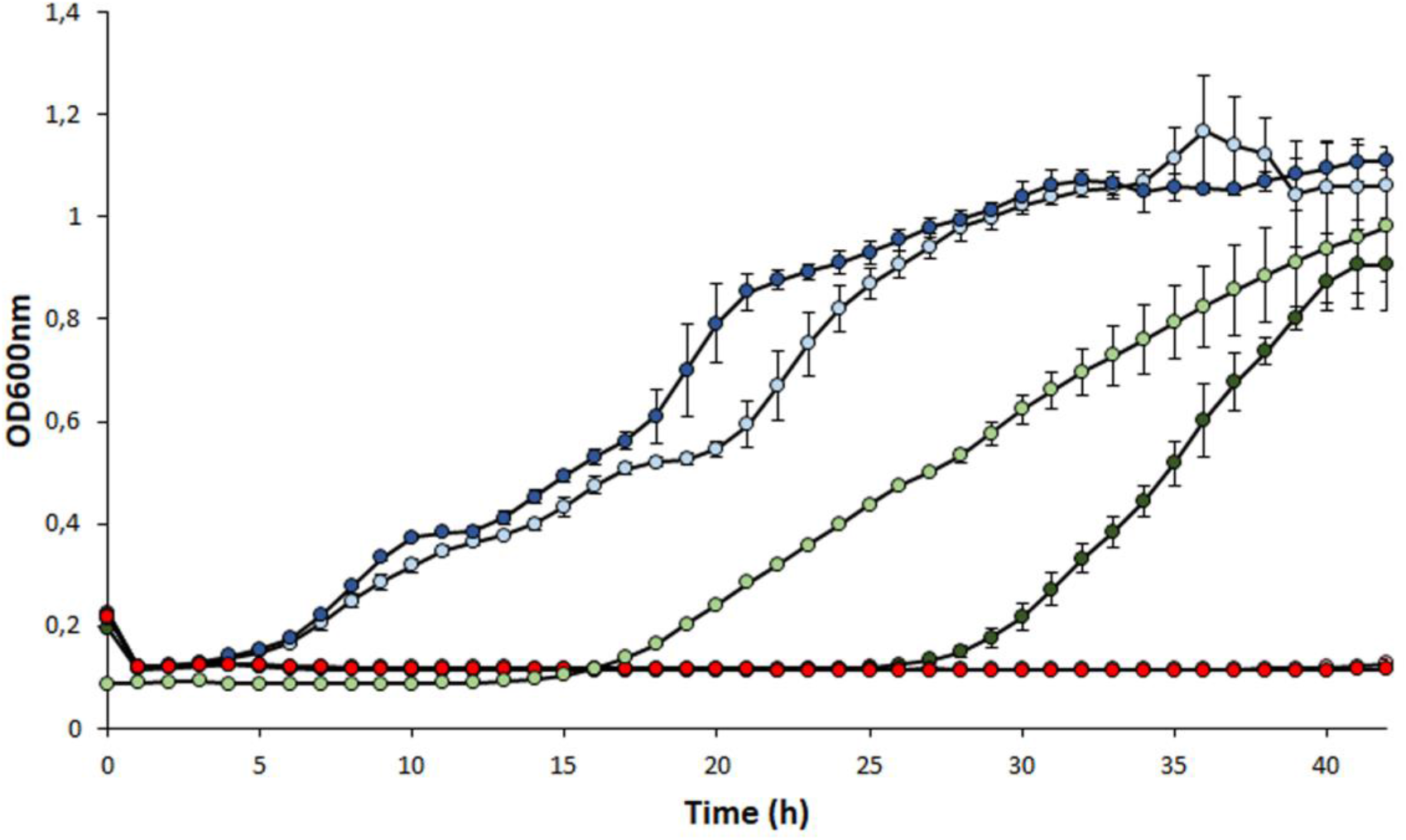
Effect of atropine in the growth of *P. aeruginosa* in presence of ciprofloxacin. Growth curves of *P. aeruginosa* PAO1 (green), *nfxB\** (blue) and of *nfxB*ΔmexCD* (red) in LB medium containing 0.25 μg/ml ciprofloxacin, with (light) or without (dark) 2 mg/ml of atropine. As shown, the wild-type *P. aeruginosa* strain grows better in presence of ciprofloxacin when atropine is added. Each OD_600nm_ represented value is the average of three biological replicates.

**Table 5.**
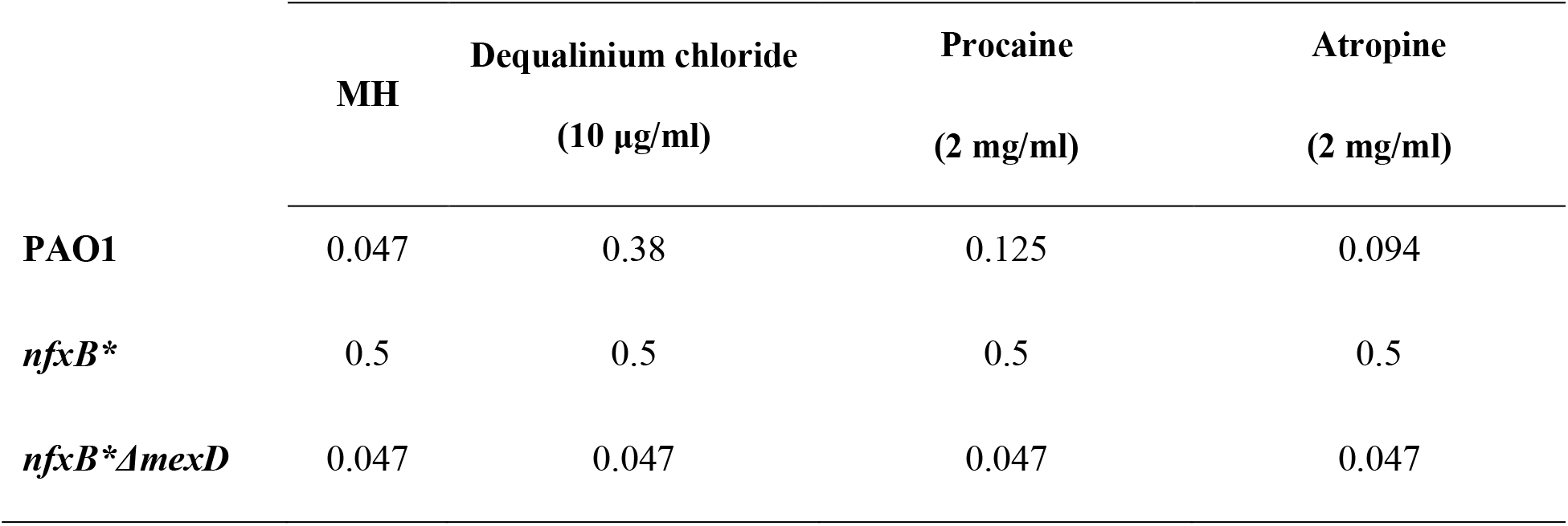
MIC (μg/ml) values of *P. aeruginosa* to ciprofloxacin, in presence or absence of inducer compounds

In order to further confirm the specific induction of *mexCD-oprJ* by these compounds, the MICs of antibiotics which are not substrates of MexCD-OprJ (tobramycin, fosfomycin and ceftazidime (8)), were also measured in presence and absence of the identified inducers. The MICs of those antibiotics did not change in presence of the inducers.

Altogether, these results indicate that the three analyzed compounds are able to transiently increase AR through the induction of the expression of the genes that encodes MexCD-OprJ. Because of that, the temporal coincidence of any of these compounds with MexCD-OprJ antibiotic substrates for treating patients may be a concern.

### MexCD-OprJ efflux pump extrudes procaine

It has been found that in some occasions, inducers of the expression of efflux pumps are also substrates of these AR determinants (22, 60–62). One example of this situation is the induction of MexXY-OprM in *P. aeruginosa* by aminoglycosides, which are substrates of this efflux pump (59). However, in other cases, as it happens in the case of *S. maltophilia* SmeVWX and SmeYZ efflux pumps (37), some of their inducers are not substrates of these efflux pumps.

To address the capacity of MexCD-OprJ to extrude its described inducers, the growth of the wild-type *P. aeruginosa* PAO1 and *nfxB*ΔmexD* strain were compared in the presence of a toxic concentration of each inducer (4 mg/ml of atropine, 4 mg/ml of procaine or 20 μg/ml of dequalinium chloride). No relevant differences in growth were observed in presence of dequalinium chloride or atropine in the MexD-deficient strain compared to the wild type. However, growth of the MexD-deficient strain was reduced in the presence of procaine (Figure 5) indicating that this compound, besides being an inducer, might also be a substrate of the MexCD-OprJ efflux pump.

**Figure 5.**
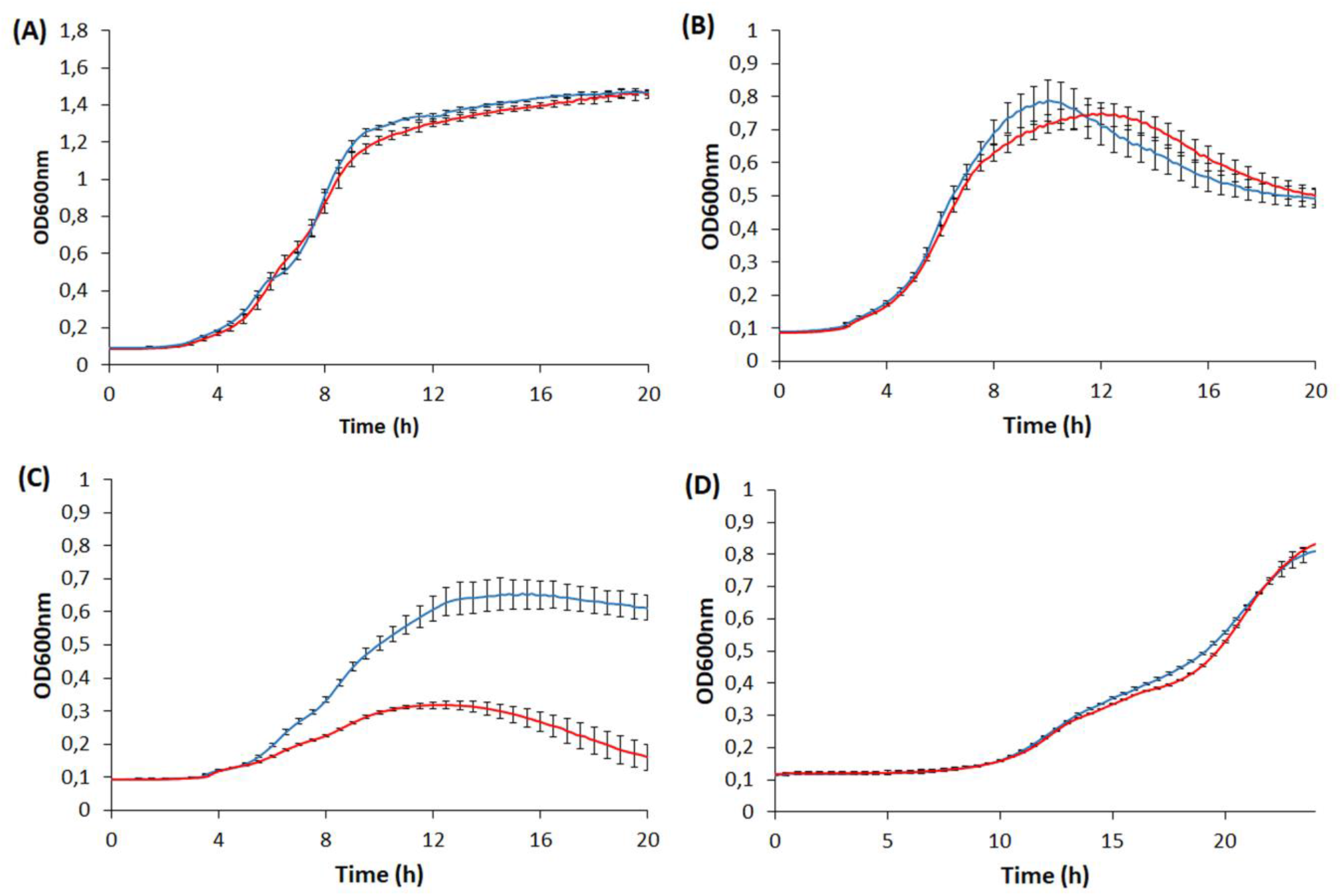
Effect of MexCD-OprJ in the susceptibility of *P. aeruginosa* to *mexCD-oprJ* inducers. PAO1 wild-type strain (blue) and *nfxB*ΔmexD* strain (red) were grown in LB as a control (A) and in presence of 4 mg/ml of atropine (B), 4 mg/ml of procaine (C) or 20 μg/ml of dequalinium chloride (D). As shown, the absence of MexCD-OprJ increases *P. aeruginosa* susceptibility to procaine. Each OD_600nm_ represented value is the average of three biological replicates.

## CONCLUDING REMARKS

The combination of Biolog plates and luminescent biosensor strains has allowed us to describe novel inducers of *P. aeruginosa* efflux pumps, some of which must be carefully taken into consideration in the clinic field; in particular, when these compounds and antibiotics are simultaneously applied. Moreover, this type of approaches, which has proven to be useful to find new inducers of efflux pumps in *P. aeruginosa*, besides informing on the putative original role of efflux pumps in non-clinical environments, may allow predicting potential conditions triggering transient AR at clinical settings.

## Acknowledgments

Work has been supported by grants from the Instituto de Salud Carlos III (Spanish Network for Research on Infectious Diseases [RD16/0016/0011]), from the Spanish Ministry of Economy, Industry and Competitivity (BIO2017-83128-R), from JPI Water StARE JPIW2013-089-C02-01) and from the Autonomous Community of Madrid (B2017/BMD-3691). The funders had no role in study design, data collection and interpretation, or the decision to submit the work for publication. PL is the recipient of a FPU fellowship, PB is the recipient of a FPI fellowship.

